# Selective inhibition of anaerobic ubiquinone biosynthesis in *Pseudomonas aeruginosa*

**DOI:** 10.64898/2026.09.23.753745

**Authors:** Julie Michaud, Yvan Caspar, Olivier Lerouxel, Tuan Anh Dinh, Philippe Simon, Murielle Lombard, Ludovic Pelosi

## Abstract

*Pseudomonas aeruginosa* is a metabolically versatile opportunistic pathogen capable of adapting to diverse and challenging environments, including the oxygen- and nutrient-limited conditions encountered during chronic infections. Its ability to utilize a broad range of carbon sources and to switch from aerobic to anaerobic respiration contributes to its persistence within the host and its resilience to therapeutic interventions. In particular, denitrification enables *P. aeruginosa* to grow under oxygen-limited conditions by using nitrate and nitrite as alternative electron acceptors, a process particularly relevant in biofilms and chronic infections such as cystic fibrosis. Ubiquinone (UQ) is essential for respiratory metabolism in *P. aeruginosa*, which possesses two distinct UQ biosynthetic pathways that ensure UQ production across a broad range of oxygen concentrations. Recent studies have identified the O_2_-independent UQ biosynthetic pathway as a key determinant of chronic infection. Because this pathway is restricted to a limited group of bacteria, we hypothesized that its selective inhibition could represent a strategy to specifically target *P. aeruginosa* under anaerobic conditions. To test this hypothesis, we screened analogues of 4-hydroxybenzoic acid (4-HB), the earliest precursor of the UQ biosynthetic pathway, and identified 4-aminobenzoic acid, also known as para-aminobenzoic acid (pABA), as a potential inhibitor. pABA primarily affects the anaerobic metabolism of *P. aeruginosa* by acting as a competing substrate for UQ biosynthetic enzymes, thereby interfering with UQ biosynthesis. This inhibitory effect positions pABA as a valuable tool for investigating, and potentially targeting, anaerobic metabolism in *P. aeruginosa*.

**Importance:** *Pseudomonas aeruginosa* is a metabolically versatile bacterium that thrives in diverse environments, including clinical settings, where it poses a major threat due to its antibiotic resistance and adaptability. Its ability to survive under both oxygen-rich and oxygen-limited conditions, through processes such as denitrification, facilitates persistence during chronic infections such as cystic fibrosis. Central to its respiratory metabolism is ubiquinone (UQ), which is essential under all oxygen conditions. *P. aeruginosa* possesses two UQ biosynthetic pathways: an O_2_-dependent and an O_2_-independent one, the latter being crucial for anaerobic survival. This pathway, widespread among *Pseudomonadota*, is associated with the maintenance of chronic infection. Thus, targeting this O_2_-independent UQ pathway represents a promising strategy for novel antimicrobial development. In this context, we have identified para-aminobenzoic acid as a potential inhibitor.

## Introduction

*Pseudomonas aeruginosa* is a versatile, Gram-negative bacterium widely found in soil, water, and hospital environments. Known for its remarkable metabolic adaptability, it can survive under diverse and often harsh conditions. This bacterium is an opportunistic pathogen, frequently causing infections in immunocompromised individuals, such as those with severe cystic fibrosis (CF), burns, or indwelling medical devices (1, 2). *P. aeruginosa* is also a major concern in clinical settings due to its intrinsic resistance to many antibiotics and its ability to acquire additional resistance mechanisms (3), making infections difficult to treat. Although *P. aeruginosa* is primarily an aerobic organism, it can withstand oxygen-limited conditions by shifting to anaerobic respiration (4, 5). Beyond its respiratory flexibility, *P. aeruginosa* can also metabolize a wide spectrum of carbon sources (6). Such metabolic versatility not only facilitates its adaptation to fluctuating nutrient conditions and colonization of diverse ecological niches but also contributes to its persistence in the human host and its resilience against therapeutic interventions.

Denitrification in *P. aeruginosa* is a key anaerobic respiratory process that allows the bacterium to survive and grow under oxygen-limited conditions. In this pathway, nitrate and nitrite serve as alternative electron acceptors in the absence of oxygen and are sequentially reduced to nitric oxide, nitrous oxide, and ultimately gaseous dinitrogen (7). This process is tightly regulated and becomes particularly important in hypoxic environments such as biofilms, where oxygen penetration is restricted. Denitrification not only supports bacterial energy production but also contributes to virulence and persistence within the host, making it an important factor in chronic infections such as CF (8, 9). The proton-motive force generated during the respiratory process of *P. aeruginosa*, i.e., arising from the transfer of electrons and protons from reduced donors to oxidized acceptors, depends exclusively on ubiquinone-9 (UQ_9_) regardless of the oxic conditions (10, 11). Indeed, like many other bacteria such as *Escherichia coli*, *P. aeruginosa* possesses two UQ biosynthetic pathways, enabling UQ production across the entire range of O_2_ concentrations (11).

As shown in Figure 1, six proteins (UbiA, UbiB, UbiD, UbiE, UbiG, and UbiX), which catalyze the prenylation, decarboxylation, and methylation of the phenyl ring of the 4-hydroxybenzoic acid (4-HB) precursor, are common to both UQ pathways (11, 12). The classical one requires O_2_ for the three hydroxylation steps (13). In *P. aeruginosa*, the flavin-dependent monooxygenases (FMOs) UbiI and UbiH, together with the di-iron hydroxylase Coq7, catalyze the O_2_-dependent hydroxylation steps, whereas in *E. coli*, the FMO UbiF fulfills the role of Coq7 (10). In *E. coli*, UbiE to UbiI and the accessory UbiK and UbiJ proteins forms a soluble complex (14). As we demonstrated previously, the O_2_-independent pathway of UQ relies on specialized enzymes, including UbiU and UbiV, which substitute for O_2_-dependent hydroxylases in the canonical UQ biosynthetic pathway found in oxic conditions, as well as the accessory protein UbiT, which is homologous to UbiJ (11, 12) (Figure 1). We also identified prephenate, an organic compound and intermediate of the aromatic amino acid pathway, as the oxygen donor for the three hydroxylation steps in the O₂-independent pathway (15).

**Figure 1:**
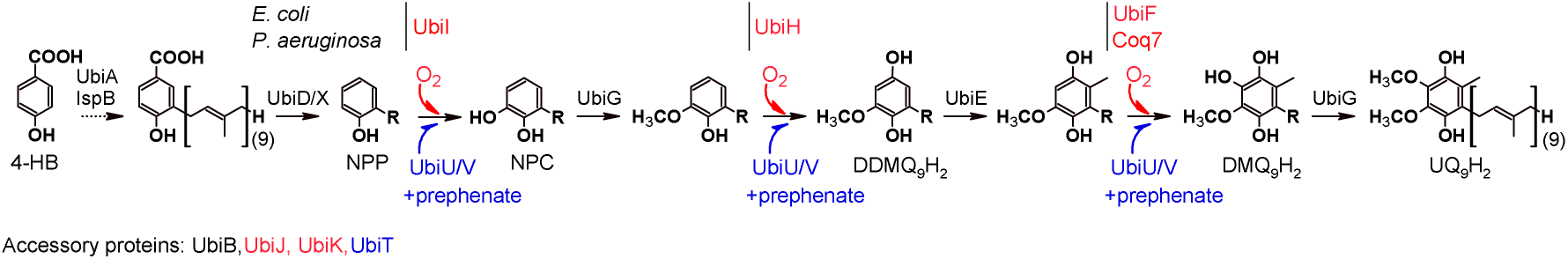
UQ_9_ biosynthesis pathway of *P. aeruginosa*. Proteins specific to the O_2_-dependent and O_2_-independent pathways are represented in red and blue, respectively. In *E. coli*, the flavin-dependent monooxygenase (FMO) UbiF fulfills a role analogous to that of Coq7. Prephenate, an intermediate in the aromatic amino acid biosynthetic pathway, serves as the oxygen donor for the three hydroxylation steps of the O₂-independent pathway. The molecules are represented in their reduced form and the nonaprenyl tail is denoted as “R”. Abbreviations: 4-HB, 4-hydroxybenzoic acid; NPP, nonaprenylphenol; NPC, nonaprenylcatechol (or 2-nonaprenyl-6-hydroxyphenol); DDMQ_9_H_2_, demethyl-demethoxy UQ_9_H_2_; DMQ_9_H_2_, demethoxy-UQ_9_H_2_; DQ_9_H_2_, demethyl-UQ_9_H_2_; UQ_9_H_2,_ ubiquinol 9.

Regarding the molecular basis of the chronic-to-acute switch in *P. aeruginosa*, recent findings show that the O_2_-independent UQ biosynthetic pathway acts as a primary determinant in sustaining chronic infection (16). As the O_2_-independent UQ biosynthetic pathway is distributed only within the *Pseudomonadota* (17), and a few species of the *Desulfobacterota* phylum (18), we propose in this study to develop a strategy for the selective inhibition of UQ biosynthesis under anaerobic conditions and to assess its potential as a drug target.

## Results

### pABA is a promising candidate for inhibiting anaerobic UQ biosynthesis in

*P. aeruginosa*. As each UQ biosynthetic pathway relies on a distinct set of hydroxylases and accessory proteins - UbiI, UbiH, Coq7, UbiK and UbiJ under aerobic conditions (10), and UbiU, UbiV and UbiT under anaerobic conditions (11) - we hypothesized that this difference could be exploited in our approach. Thus, we tested the ability of 4-HB analogues to inhibit the UQ_9_ biosynthetic pathway in *P. aeruginosa* PAO1 under anaerobic conditions. First, we screened analogues modified at the para position of the carboxylate group. Each compound, dissolved in DMSO, was tested at a final concentration of 1 mM in minimal culture medium containing succinate as the respiratory carbon source and supplemented with KNO_3_ as the terminal electron acceptor. The endogenous UQ_9_ content of bacterial cells was measured by high-performance liquid chromatography (HPLC) with electrochemical detection (ECD) coupled to mass spectrometry (MS) and compared to a control condition in which only DMSO was added. As shown in Figure 2, 4-fluorobenzoic acid and 4-methylbenzoic acid had no significant effect on UQ_9_ content, while 4-mercaptobenzoic acid and, surprisingly, 4-HB led to slight decreases of 24% and 17%, respectively. A stronger inhibition was observed with 4-nitrobenzoic acid (51%). Ultimately, the most potent inhibitor was 4-aminobenzoic acid (pABA), which strongly reduced UQ_9_ biosynthesis by 90% compared to the control condition (Figure 2). Based on these results, we then refocused our screen on commercially available pABA analogues bearing a hydroxyl, methoxy, methyl, or amino group at the ortho or meta position of the carboxylate group (Figure S1). All compounds, tested at a final concentration of 1 mM, led to a reduction in UQ_9_ content ranging from 19% to 46% compared to the control condition. Notably, pABA still produced the strongest inhibitory effect (Figure S1) and was therefore selected for further use in this study.

**Figure 2:**
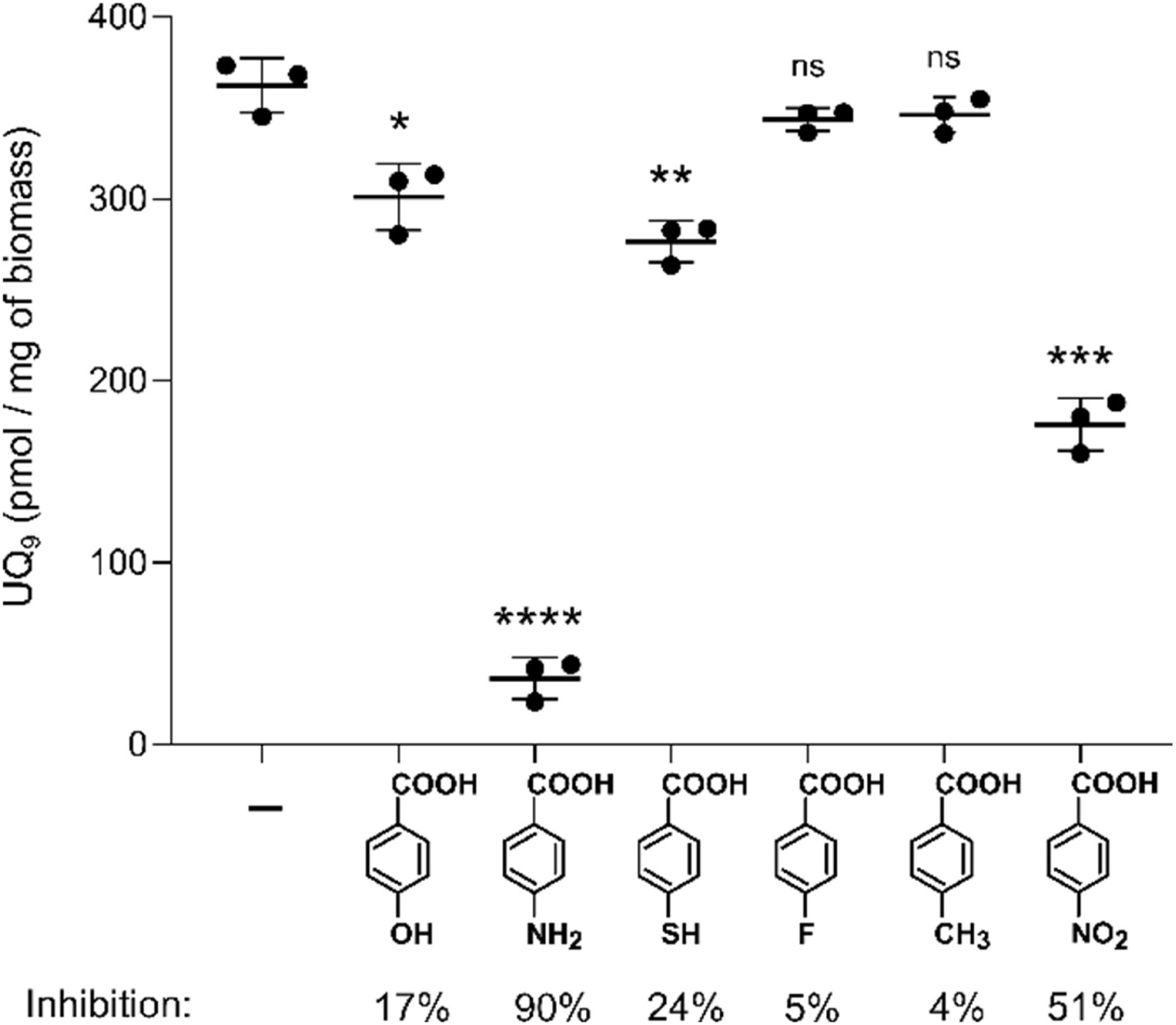
Ability of 4-HB analogues to inhibit UQ_9_ biosynthesis in *P. aeruginosa* PAO1 under anaerobic conditions. Each 4-HB derivative, dissolved in DMSO, was tested at a final concentration of 1 mM in minimal culture medium containing succinate and supplemented with KNO₃. UQ₉ levels of PAO1 were quantified and normalized to biomass (n=3). Statistical significance was determined using an unpaired Student’s t-test compared to the untreated control, which contained only DMSO (-). \**P* < 0.05; \*\**P* < 0.01; \*\*\**P* < 0.001; \*\*\*\**P* < 0.0001; ns, not significant. The structure of each compound is shown. From left to right: 4-hydroxybenzoic acid, 4-aminobenzoic acid, 4-mercaptobenzoic acid, 4-fluorobenzoic acid, para-toluic acid and 4-nitrobenzoic acid.

**Figure 3:**
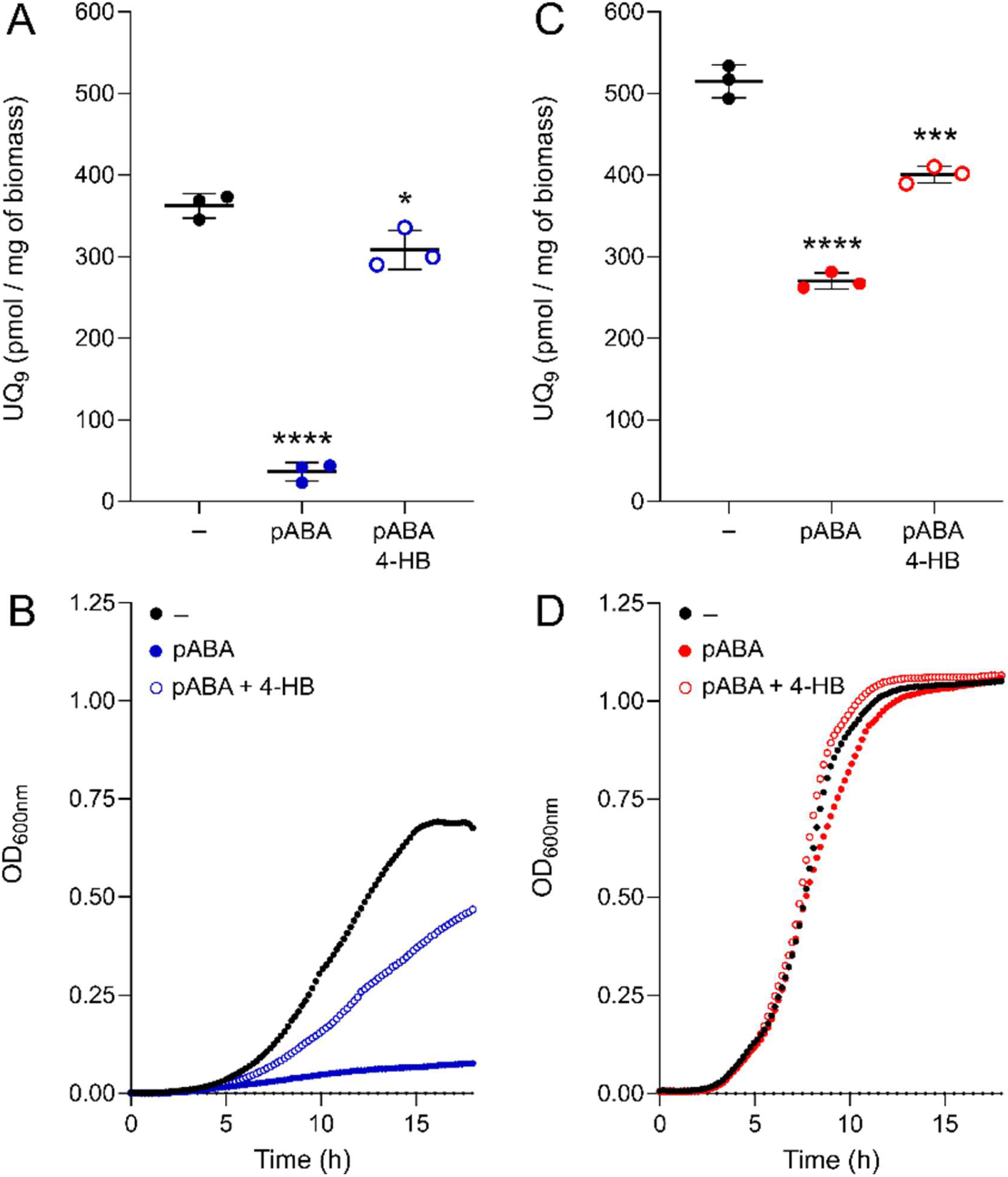
pABA inhibits anaerobic UQ_9_ biosynthesis and bacterial growth in *P. aeruginosa* PAO1. Quantification of UQ_9_ extracted from PAO1 cultures grown under anaerobic **(A)** or aerobic **(C)** conditions in the presence of 1 mM pABA alone or in combination with 1 mM 4-HB (n=3). Corresponding growth curves under anaerobic **(B)** and aerobic **(D)** conditions were obtained by monitoring OD_600_ at 37 °C (n=7). Statistical significance was determined using an unpaired Student’s t-test, comparing each condition with the untreated control, which contained only DMSO (–). \**P* < 0.05; \*\*\**P* < 0.001; \*\*\*\**P* < 0.0001.

**Figure 4:**
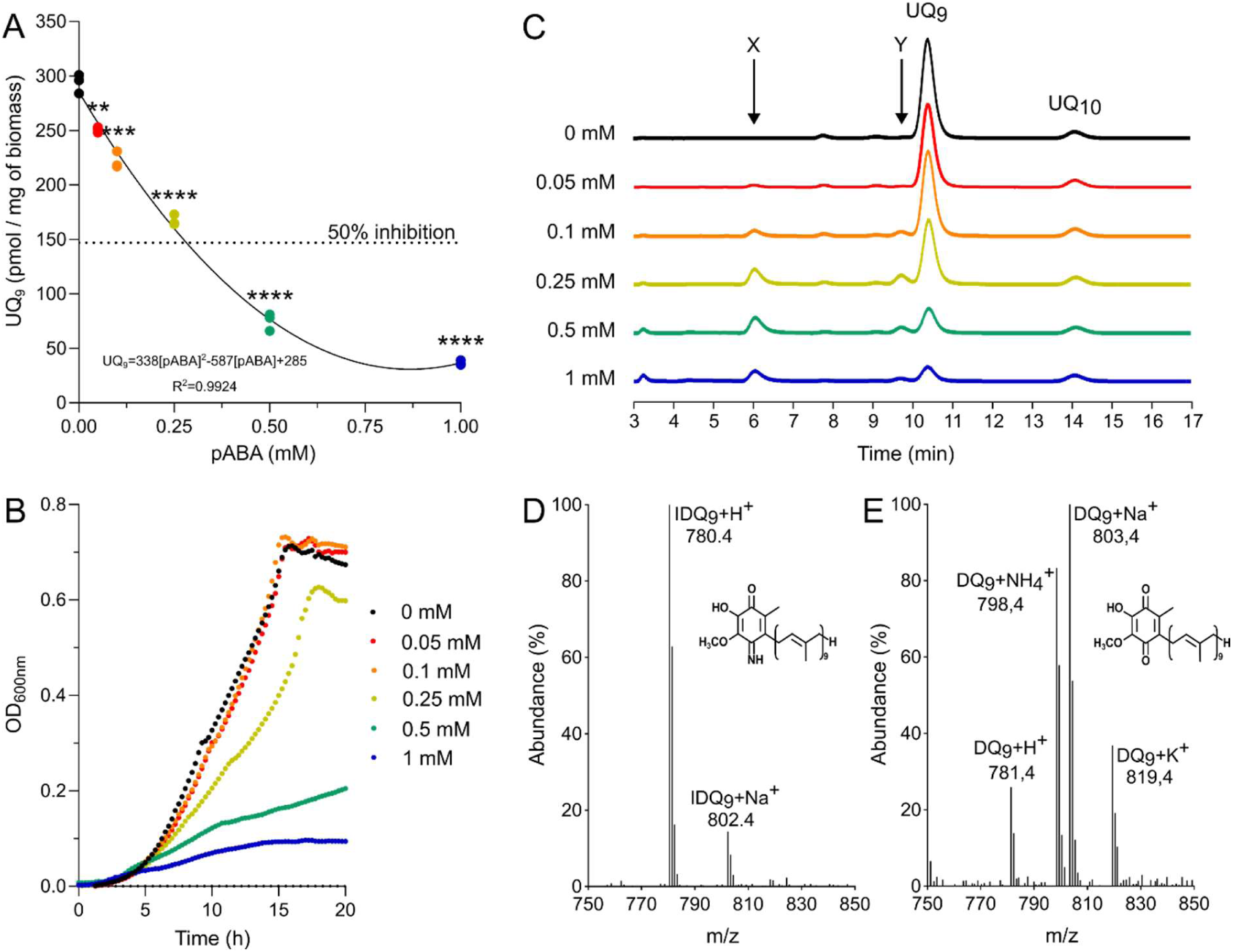
pABA competitively inhibits anaerobic UQ_9_ biosynthesis in *P. aeruginosa* and affects virtually several steps of the pathway. **(A)** Quantification of UQ_9_ extracted from *P. aeruginosa* cultures grown under anaerobic conditions in the presence of increasing concentrations of pABA and mathematical modeling of the dose-response relationship between pABA concentration and UQ_9_ content. Data were fitted with a quadratic regression, whose equation is shown on the graph, reflecting a non-linear decrease in UQ_9_ content with increasing pABA concentration. The dashed horizontal line indicates the 50% inhibition threshold relative to the control condition. **(B)** Corresponding growth curves (see legend of Figure 3), and **(C)** HPLC-ECD analysis of the corresponding lipid extracts. The chromatograms are representative of results from three independent experiments. The peaks corresponding to UQ_8_ and the UQ_10_ standard are indicated. Compound X and Y eluting at 6.2 and 9,7 min are marked. Mass spectra of compound X **(D)** and Y **(E)**. H^+^, NH_4+_, Na^+^, and K^+^ adducts corresponding to these molecules are indicated if they are detected. Inset, proposed structure of compound X and Y in its oxidized form. In (A), Statistical significance was determined using an unpaired Student’s t-test, with the untreated control containing only DMSO used as the reference. ****P < 0.0001; ***P < 0.001; **P < 0.01.

### pABA inhibits mainly the anaerobic UQ biosynthesis in *P. aeruginosa*, impairing bacterial growth

We investigated the ability of pABA, tested at a final concentration of 1 mM, to inhibit UQ_9_ biosynthesis in *P. aeruginosa* PAO1 across the entire O₂ range, and consequently its effect on bacterial growth. As expected, concomitant with the strong decrease in UQ_9_ content previously observed under anaerobic conditions in the presence of pABA, *P. aeruginosa* growth was strongly impaired (Figures 3A and 3B). Control experiments also showed that the addition of 4-HB to the growth medium largely counteracted the negative effects of pABA, restoring both UQ_9_ biosynthesis and bacterial growth, albeit partially (Figures 3A and 3B). In contrast, adding pABA to the culture medium under ambient air led to a moderate reduction in UQ_9_ content, decreasing it by only 47% compared to the control, while bacterial growth remained unaffected (Figures 3C and 3D). Together, these findings demonstrate that pABA inhibits the UQ_9_ biosynthetic pathway with significantly greater potency under anaerobic than aerobic conditions, most likely acting as a competitive inhibitor, as shown below.

### pABA acts as a competing substrate for UQ_9_ biosynthetic enzymes, while influencing virtually every step of the pathway

We then examined how and to what extent pABA could affect UQ_9_ biosynthesis in *P. aeruginosa* PAO1. Bacteria were cultured under anaerobic conditions in the presence of pABA (50 µM to 1 mM, final concentration). The endogenous UQ_9_ content was measured in bacterial cells and compared to the control condition. Figure 4A shows that the UQ_9_ content decreased with increasing concentrations of pABA in the medium, with 0.28 mM (deduced from mathematic model) yielding an ∼50% decrease compared to the untreated condition. Concomitantly, the growth of *P. aeruginosa* was impaired from 0.25 mM pABA onward (Figure 4B). Treatment with pABA caused the accumulation of a redox compound that eluted at around 6 min (Figure 4C, compound X). Mass spectrometry (MS) analysis of this peak showed a predominant proton adduct (M+H⁺) at m/z 780.4, together with a minor adduct (M+Na⁺) at m/z 802.4 (Figure 4D). The two species are compatible with a monoisotopic mass of 779.4 g·mol^-^¹, which most likely corresponds to that of 4-imino-6-demethyl-UQ_9_, hereafter referred to as IDQ_9_ (Figure 4D, inset). IDQ_9_ can be reduced to 4-amino-6-demethyl-UQ_9_ (IDQ_9_H_2_). Concomitantly, another redox compound, eluting at around 9.7 min, also accumulated (Figure 4C, compound Y). MS analysis of this peak showed a monoisotopic mass of 780.4, corresponding to 6-demethyl-UQ_9_ (DQ_9_) (Figure 4E, inset), which can be reduced to 6-demethyl-UQ_9_H_2_. This analogue, which corresponds to the 4-HB-derived counterpart of IDQ_9_H_2_ (Figure 1, compound DQ_9_H_2_), is the natural product of UbiU and UbiV at the third hydroxylation step and the second natural substrate of UbiG after nonaprenylcatechol (Figure 1, compound NPC).

To trace the origin of newly synthesized IDQ_9_, ^13^C_6_-pABA (250 µM final concentration) was added to anaerobic cultures, and the quinone content together with the isotopic labeling pattern was analyzed by HPLC-ECD-MS. The accumulated compound eluting at 6 min showed a MS signal for the proton adduct (M+H⁺) at m/z 786.4, corresponding to the expected mass of ^13^C_6_-IDQ_9_ (Figure 5A). Integration of the signal at m/z 786.4 showed the absence of the previously accumulated ¹²C₆-IDQ_9_ (Figure 5B). This result showed that IDQ_9_ originates exclusively from the exogenously supplied ^13^C_6_-pABA. In parallel, incubations with ^13^C_6_-pABA produced no detectable ^13^C_6_-UQ_9_, suggesting that pABA does not serve as a ring precursor for UQ biosynthesis in *P. aeruginosa* (Figures 5C and 5D). Thus, according to the sequence of reactions proposed in the literature for the UQ_9_ biosynthetic pathway (11), the formation of IDQ_9_H_2_ would result from the prenylation of added pABA, decarboxylation at C1, hydroxylation and then methylation at C5, hydroxylation at C1, methylation at C3, and finally hydroxylation at C6 (Figure 5E). Thus, IDQ_9_H_2_, which is not methylated by UbiG, appears to be the “dead-end” product of the UQ_9_ pathway in *P. aeruginosa* cells treated with pABA (Figure 5E). Collectively, these results unequivocally demonstrate that pABA primarily affects the anaerobic metabolism of *P. aeruginosa* by acting as a competing substrate for UQ_9_ biosynthetic enzymes, thereby interfering with UQ_9_ biosynthesis (Figure 5E). This suggests an extensive mechanism of action affecting nearly every step of the pathway, positioning pABA as a valuable tool for studying, and potentially controlling, anaerobic metabolism in this opportunistic pathogen.

**Figure 5:**
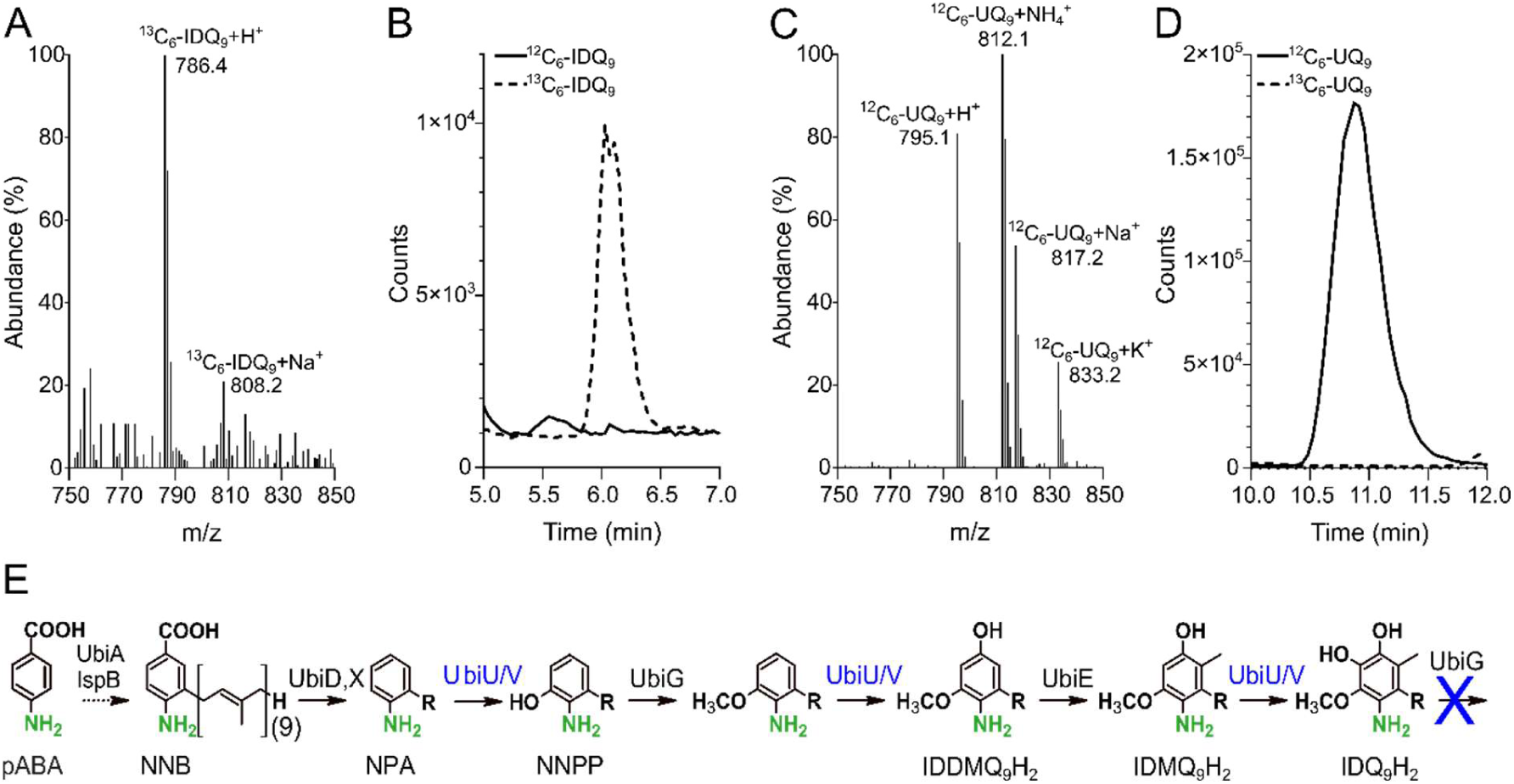
Added pABA serves as the primary source of IDQ₉ but does not act as a precursor for the UQ ring in *P. aeruginosa*. Mass spectrum of synthetized ^13^C6-IDQ_9_ **(A)** and ^12^C6-UQ_9_ **(B)**. H^+^, NH_4+_, Na^+^, and K^+^ adducts corresponding to these molecules are indicated when detected. **(C-D)** HPLC-MS profiles for SIM with H^+^ adduct (IDQ_9_+H^+^ and UQ_9_+H^+^) corresponding to ^12^C6-IDQ_9_ or ^12^C6-UQ_9_ shifted by +6 mass units (^13^C6-IDQ_9_ or ^13^C6-UQ_9_). **(E)** Sequence of reactions proposed for UQ_9_ biosynthesis in *P. aeruginosa* in presence of pABA as precursor. The molecules are represented in their reduced form and the nonaprenyl tail is denoted as “R”. Abbreviations: NNB, 3-nonaprenyl-4-aminobenzoic acid; NPA, 2-nonaprenyl-aniline; NNPP, 2-amino-3-nonaprenylphenol; IDDMQ_9_H_2_, imino-demethyl-demethoxy-UQ_9_H_2_; IDMQ_9_H_2_, imino-demethoxy-UQ_9_H_2_; and IDQ9H_2_, imino-demethyl-UQ_9_H_2_.

### UbiG may not be able to methylate IDQ_9_H_2_

The accumulation of IDQ_9_ could suggest a defect in either the UbiU-UbiV hydroxylase, resulting in a failure to release the product from the enzyme, and/or in UbiG, which would be unable to catalyze the final methylation step despite recognizing and modifying 2-amino-3-nonaprenylphenol (Figure 5E, compound NNPP). Since the three-dimensional (3D) structure of the UbiU-UbiV heterodimer is not yet known, and the hydroxylation mechanism associated with this novel family of hydroxylases remains poorly understood, we considered structural modeling of this hypothesis premature given the current structural and mechanistic uncertainties. In contrast, the crystal structure of *E. coli* UbiG in complex with S-adenosyl-L-homocysteine, an analogue of its co-substrate S-adenosyl-L-methionine (SAM), has been determined (19, 20). Thus, using AlphaFold3, we generated a high-fidelity 3D model of *P. aeruginosa* UbiG in complex with SAM and subsequently docked either its natural substrate, DQ_9_H_2_ (Figure 6A), or the alternative substrate, IDQ_9_H_2_ (Figure 6B), into the active-site cavity. Both substrates are stabilized by residues Asp124, His125, and His177. As illustrated in the overlapped models, the orientation of the aromatic ring differs substantially between the two substrates, resulting in marked changes in the predicted hydrogen-bonding interactions that favor stronger binding of the natural substrate (Figures 6C and 6D). Moreover, the distance between the oxygen atom at C6 of the redox moiety and the carbon atom of the methyl group transferred from SAM is significantly increased with IDQ_9_H_2_ (3.53 vs 3.01 Å) (Figure 6D).

**Figure 6.**
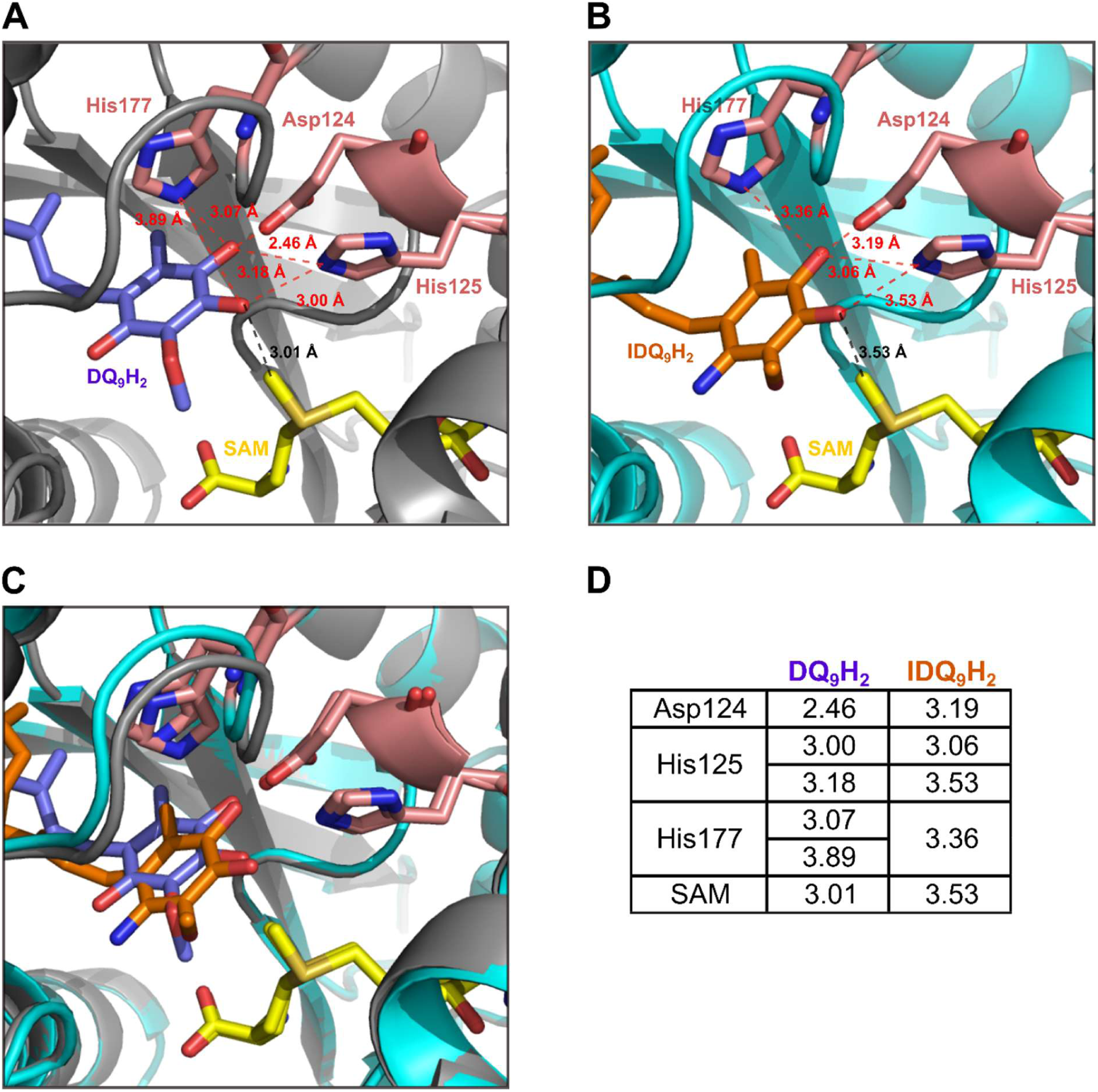
Three-dimensional model of UbiG from *P. aeruginosa* in complex with SAM and docked DQ_9_H_2_ or IDQ_9_H_2_. **(A)** AlphaFold-predicted structure of UbiG showing the docking pose of DQ_9_H_2_ in the presence of SAM (ipTM = 0.9096, pTM = 0.9366). **(B)** Docking pose of IDQ_9_H_2_ in the same binding cavity in the presence of SAM (ipTM = 0.9180, pTM = 0.9407). **(C)** Superposition of the DQ_9_H_2_- and IDQ_9_H_2_-bound models. **(D)** Distances (≤4Å) between the docked ligands, SAM, and the residues lining the binding cavity (Asp124, His125, and His177). Dashed lines indicate the distances measured between the corresponding atoms. DQ_9_H_2_: demethyl-UQ_9_H_2_; IDQ_9_H_2_: imino-demethyl-UQ_9_H_2_.

These results were compared to those obtained with NPC and NNPP, both of which are recognized and modified by UbiG (Figures 1 and 5E). The docking results revealed no substantial differences between the simulations performed with NPC and NNPP as ligands (Figures S2A and S2B). In particular, the predicted hydrogen-bonding networks were largely conserved between the two complexes, providing no clear structural evidence for preferential binding of either substrate, nor for differential positioning that would favor methylation of one substrate over the other (Figures S2C and S2D). Although this *in silico* approach remains preliminary, the results obtained suggest that UbiG may be unable to methylate IDQ_9_H_2_ because of a potentially unfavorable positioning of the redox moiety. Nevertheless, further biochemical data will be required to substantiate this hypothesis.

### *P. aeruginosa* pathogenic strains are sensitive to pABA mainly in anaerobiosis

To generalize the effect of pABA on the anaerobic metabolism of *P. aeruginosa*, we screened multiple strains isolated from blood cultures, as well as from intensive care patients with hospital-acquired pneumonia or from CF patients. Three strains were tested for each type of isolate. In all cases, the addition of 1 mM pABA to the culture medium strongly impaired anaerobic growth (Figure 7A), concomitant with a drastic reduction in bacterial UQ_9_ content, ranging from ∼87 to 92% compared to the control condition (Figure 7B). As previously observed for the strain PAO1, the UQ_9_ content of pathogenic strains was significantly affected by pABA under ambient air (∼38 to 51% inhibition), but not enough to generate a bacterial growth deficit (Figures S3A and S3B). Together, these results demonstrate that the inhibitory effect of pABA on UQ_9_ biosynthesis and anaerobic growth is not restricted to the laboratory strain PAO1 but is a conserved feature across clinically relevant *P. aeruginosa* isolates, regardless of their origin.

**Figure 7:**
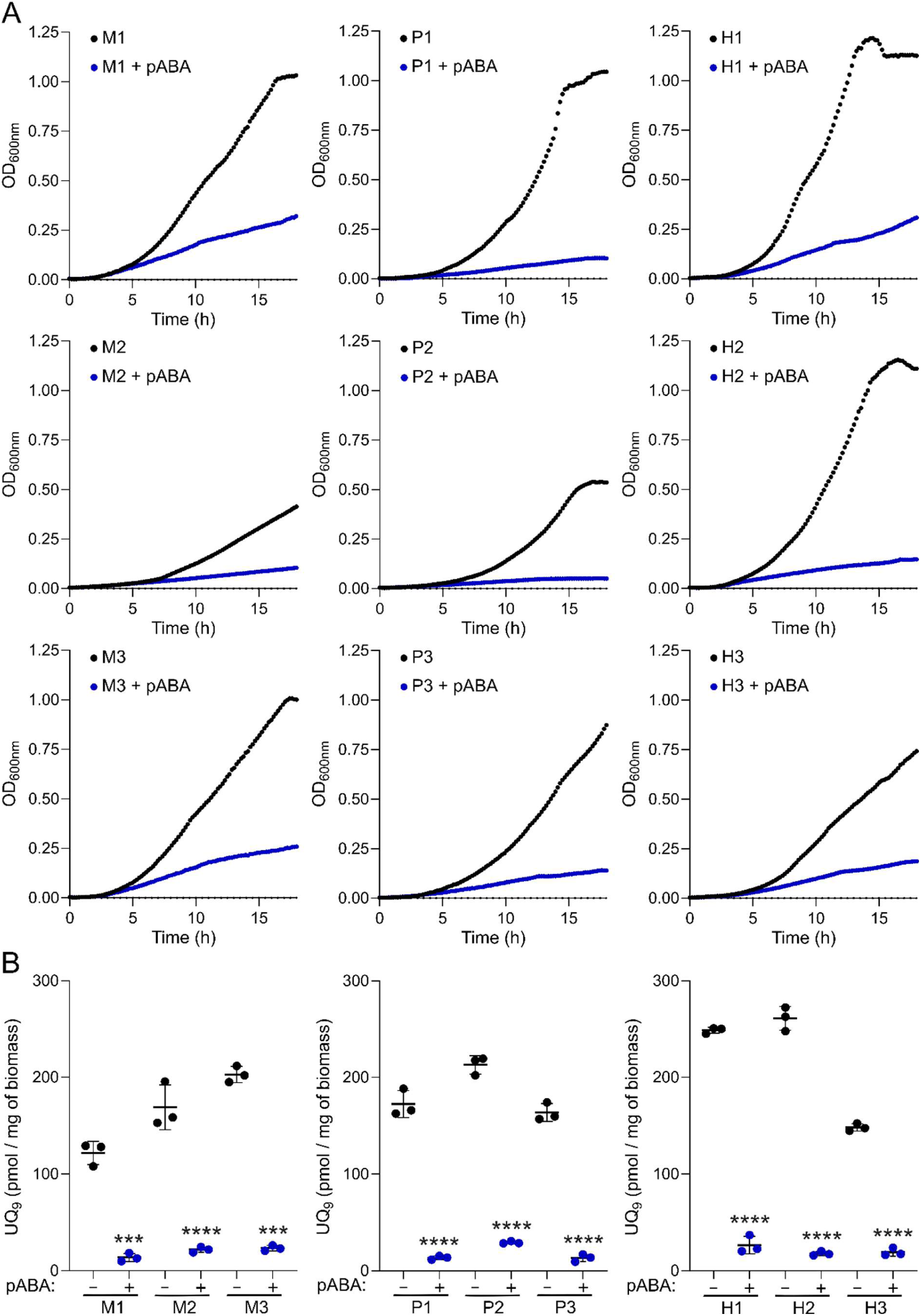
Effect of pABA on clinical strain UQ_9_ contents and growth under anaerobic conditions. Clinical strains isolated from patients (M1 to M3 from cystic fibrosis patients’ sputum; P1 to P3 from intensive care patients’ tracheal aspirate and H1 to H3 from blood culture) were grown anaerobically in minimal medium supplemented with 1 mM pABA. **(A)** Growth curves were obtained by monitoring OD₆₀₀ over 18 h at 37 °C (n=7). **(B)** UQ_9_ levels were quantified and normalized to biomass (n=3). Statistical significance was determined using an unpaired Student’s t-test, with the untreated control containing only DMSO (-) used as the reference. \*\*\**P* < 0.001; \*\*\*\**P* < 0.0001.

### pABA does not interact with antibiotics commonly used to fight *P. aerugino*s*a* infection

To investigate potential synergistic effects between pABA and antibiotic classes commonly used for *P. aeruginosa* infections (namely fluoroquinolones, β-lactams, and aminoglycosides), synergy testing with ciprofloxacin, meropenem, and gentamicin was performed using a checkerboard assay. As this method is primarily used under aerobic conditions, the minimum inhibitory concentrations (MICs) of the three antibiotics were first determined under anaerobic conditions (Table S1). The calculated mean fractional inhibitory concentration index (FICI) confirmed that no synergistic effect was observed, as the addition of pABA did not reduce the MIC of any of the three antibiotics and showed that pABA did not antagonize the activity of any of the tested antibiotics either (Table S1).

## Discussion

*P. aeruginosa* exhibits remarkable metabolic flexibility, particularly through its ability to switch from aerobic to anaerobic respiration, which contributes to its persistence and adaptability within the host (3). Under oxygen-limited conditions, the bacterium can rely on denitrification to sustain energy production and growth. This respiratory adaptation is particularly important in oxygen-depleted environments such as biofilms of patients with chronic CF infections, where it promotes bacterial survival and may contribute to reduced susceptibility to antimicrobial therapies (7). This observation is consistent with our results, which showed increased MICs for gentamicin and meropenem under anaerobic conditions (Table S1). A few years ago, we demonstrated that UQ_9_ was essential for *P. aeruginosa* survival under anaerobic conditions (11). More recently, Cao *et al*. showed that UQ is a key determinant of bacterial persistence during chronic infection, highlighting the O_2_-independent UQ_9_ biosynthetic pathway as a promising therapeutic target (16). To investigate this hypothesis, we chose to use 4-HB analogues, as this approach had already been shown to effectively inhibit UQ synthesis in the aerobic bacterium *Francisella novicida*, the causative agent of tularemia (21).

In our study, we identified pABA as an inhibitor of anaerobic UQ_9_ biosynthesis, strongly and selectively impairing the anaerobic growth of *P. aeruginosa* strains of both laboratory and clinical origin. pABA is an analogue of 4-HB in which the para-hydroxyl group is replaced by an amino group. Among the analogues tested, pABA was the most effective inhibitor of UQ_9_ biosynthesis under anaerobic conditions, highlighting the critical role of the para-amino functionality in this activity. Notably, introducing additional substituents onto the aromatic ring did not enhance the inhibitory activity of pABA derivatives. We showed that pABA does not serve as a ring precursor for UQ biosynthesis in *P. aeruginosa*, as previously observed in *E. coli*, the plant *Arabidopsis*, or mammalian cells (22, 23). So far, pABA has been reported to be a UQ precursor only in *S. cerevisiae* (24). In *E. coli*, prenylated pABA is decarboxylated and then hydroxylated to yield 2-amino-3-octaprenylphenol, which has been suggested to be a “dead-end” product (23). Thus, the modification of the aromatic ring of pABA appears to be prematurely halted in this bacterium after the action of UbiI, whereas the accumulation of IDQ_9_ in *P. aeruginosa* cells suggests that pABA may act as a competing substrate for all Ubi enzymes, thereby interfering with multiple steps of the O_2_-independent UQ_9_ biosynthetic pathway. In mammalian cell cultures, pABA decreases UQ levels (23, 25). However, this compound is rapidly and efficiently eliminated from the human body (26), limiting the risk of interference with UQ biosynthesis. Thus, pABA appears to interfere with mammalian UQ biosynthesis only in cell culture (27).

We found that pABA-mediated inhibition preferentially targets the anaerobic UQ biosynthetic pathway of *P. aeruginosa*. This specificity is particularly relevant in the context of chronic pulmonary infections in CF patients, for whom *P. aeruginosa* is a major cause of morbidity and mortality (28). The transition of *P. aeruginosa* from acute to chronic infection has been associated with upregulation of the O_2_-independent UQ biosynthetic pathway, which contributes to bacterial survival and growth within biofilms (16). Here, we showed that pABA inhibits the anaerobic growth of clinical isolates obtained from CF patients with chronic *P. aeruginosa* infections, suggesting that pABA remains effective against strains already adapted to biofilm-associated survival. Importantly, pABA did not interfere with the activity of commonly used antibiotics. Taken together, these findings highlight the potential of pABA as a promising therapeutic candidate for the treatment of chronic *P. aeruginosa* infections.

The differential inhibition of the aerobic and anaerobic pathways raises questions regarding the underlying mechanism, as 4-HB serves as a common precursor for both biosynthetic routes (11, 12). We know that pABA competitively inhibits the incorporation of endogenous 4-HB into the UQ biosynthetic pathway. One possible explanation for our results is that intracellular 4-HB availability differs between aerobic and anaerobic conditions, thereby affecting the sensitivity of the two pathways to pABA. In addition, *P. aeruginosa* possesses a broad repertoire of multidrug efflux pumps that actively extrude a wide range of compounds, thereby reducing the intracellular accumulation and efficacy of numerous antibiotics (29). One possibility is that pABA accumulates to higher levels under anaerobic conditions because reduced efflux activity limits its export from the cell. However, this hypothesis appears inconsistent with previous reports showing that hypoxia enhances antibiotic resistance in *P. aeruginosa* through changes in the composition and activity of multidrug efflux pumps (30). Consistent with these findings, we observed experimentally that the MICs of meropenem and gentamicin were 4- and 16-fold higher, respectively, under anaerobic than under aerobic conditions (Table S1). These results therefore argue against a general increase in efflux activity under anaerobiosis and suggest that other mechanisms may contribute to the increased sensitivity of *P. aeruginosa* to pABA under this condition. Notably, the MIC of pABA determined in our study was 75 mg/L, within the same order of magnitude as that determined for gentamicin. In contrast, the MIC previously reported for *P. aeruginosa* grown under aerobic conditions in a rich medium was substantially higher (1500 mg/L) (31). This marked difference may reflect the distinct experimental conditions used in this study compared with ours, and further supports the possibility that the activity of pABA is strongly influenced by the physiological state of the bacterium.

The other hypothesis is based on the differences between the two biosynthetic pathways. Indeed, they rely on a distinct set of hydroxylases and accessory proteins: UbiI, UbiH, Coq7, UbiK, and UbiJ under aerobic conditions, and UbiU, UbiV, and UbiT under anaerobic conditions (10–12). The accumulation of IDQ_9_, and to a lesser extent DQ_9_, suggests a defect in UbiG-mediated methylation rather than in UbiU/UbiV-mediated release of the hydroxylation product. The molecular modeling results obtained in our study support this conclusion.

However, UbiG is common to both UQ_9_ biosynthetic pathways (10, 11). A previous study has shown that most Ubi proteins in *E. coli* are organized into a multienzyme complex under aerobic conditions (14) and a similar organization may also occur in *P. aeruginosa*. This raises the question of how the composition and architecture of this complex are modulated to accommodate hydroxylase sets adapted to different oxygenation conditions. Although this question remains unresolved, the differential sensitivity of the aerobic and anaerobic pathways, despite their shared use of 4-HB and UbiG, points to a potential role for the distinct enzyme composition and organization of the two biosynthetic machineries. We therefore propose that differences in the architecture or substrate-channeling properties of the anaerobic Ubi complex may contribute to the preferential incorporation of pABA under anaerobic conditions, potentially facilitating its use as an alternative substrate up to the final methylation step.

Collectively, our data identify pABA as a valuable tool for targeting anaerobic metabolism in *P. aeruginosa*. However, further biochemical and structural studies will be required to determine the precise molecular basis of this selectivity and to establish whether pABA or optimized pABA derivatives can be exploited to selectively disrupt anaerobic respiration during chronic infection.

## Material and methods

### Bacterial strains, growth conditions and chemicals

Several *P. aeruginosa* strains were used in this study, as listed in Table S2. Strains were grown on Lysogeny broth (LB) agar plates incubated at 37°C for 24 h. Aerobic cultures were performed in minimal medium composed of 8.5 g/L Na_2_HPO_4_ 2H_2_O, 3 g/L KH_2_PO_4_, 0.5 g/L NaCl, 2.46 g/L (NH_4_)_2_SO_4_, 0.8 mg/L CaCl_2_, 0.15 g/L MgCl_2_, 8.5 mg/L FeSO_4_ 7H_2_O, 1.2 mg/L Na_2_MoO_4_, 0.1 g/L Thiamine-HCl, 0.1 g/L Cysteine-HCl, 5 g/L casein hydrolysate and 4 g/L succinate. For anaerobic growth, the medium was supplemented with 10 g/L KNO_3_ and deoxygenated with argon before sterilization. Anaerobic cultures for quinone content analysis were carried out in 13 mL Hungate tubes injected with 100 µL of overnight microaerobic cultures in closed 1.5 mL microtubes filled to 1.4 mL The cultures were incubated for 20 h at 37 °C. For aerobic growth, 5 mL of medium were inoculated with 50 µL of overnight 5 mL aerobic cultures and incubated for 20 h at 37°C with shaking at 180 rpm. Growth studies in ambient air were conducted as described previously (11) in 200 µL cultures in 96-well plates inoculated with overnight-cultures to obtain a starting 600 nm optical density (OD) of 0.001. The plates were maintained at 37°C during growth. For anaerobic growth curves, the plate reader was placed into a Whitley H35 hypoxic station with a 90% N_2_, 5% H_2_, 5% CO_2_ atmosphere. OD was monitored in the hypoxic station during 18 h at 10 min intervals using a BioTek Epoch 2 microplate spectrophotometer. Analogues of 4-HB; para-aminobenzoic acid, ^13^C6 para-aminobenzoic acid, 4-mercaptobenzoic acid, 4-fluorobenzoic acid, para-toluic acid, 4-nitrobenzoic acid, 4-aminosalicylic acid, 4-amino-3-hydroxybenzoic acid, 4-amino-2-methoxybenzoic acid, 4-amino-3-methoxybenzoic acid, 4-amino-2-methylbenzoic acid, 4-amino-3-methylbenzoic acid, 2,4-diaminobenzoic acid and 3,4-diaminobenzoic acid were tested as potential inhibitors of UQ_9_ biosynthesis in *P. aeruginosa*. These analogues were dissolved in DMSO (1 mM final concentration) and added to the culture medium prior to the inoculation.

### Lipid extractions and quinone analysis

Cultures were cooled on ice for 30 minutes prior to centrifugation at 3,200 × *g* for 10 minutes at 4°C. The resulting cell pellets were washed with 1 ml of cold PBS and transferred into pre-weighed 1.5 ml microtubes. Following centrifugation at 12,000 × *g* for 1 minute at 4°C, the supernatant was removed. The microtubes were then weighed to determine the wet weight of the recovered biomass. Pellets were stored at -20°C. Quinone extraction from biomass was performed as previously described (32). Lipid extracts in ethanol, corresponding to 1 mg of wet cell mass, were analyzed by HPLC with ECD coupled to MS. Elution was carried out over 17 minutes using a BetaBasic-18 column with a flow rate of 1 mL/min. The mobile phase consisted of 50% methanol, 40% ethanol, and 10% of a mix containing 90% isopropanol, 10% 1 M ammonium acetate, and 0.1% trifluoroacetic acid. MS detection was performed on a MSQ spectrometer (Thermo Scientific) with electrospray ionization in positive mode (probe temperature 400°C, cone voltage 80V). Single ion monitoring (SIM) detected the following compounds: UQ_9_ (M^+^ NH_4+_), *m/z* 812 to 813, 9 to 13 min; UQ_10_ (M^+^ NH_4+_), *m/z* 880 to 881, 12 to 16 min; DQ_9_ (M^+^ H^+^), *m/z* 781 to 782, 9 to 13 min; [^13^C_6_]-DQ_9_ (M^+^ H^+^), *m/z* 787 to 788, 9 to 13 min; IDQ_9_ (M^+^ H^+^), *m/z* 780 to 781, 4 to 8 min; [^13^C_6_]-IDQ_9_ (M^+^ H^+^), *m/z* 786 to 787, 4 to 8 min; UQ_9_ (M^+^ H^+^), *m/z* 795 to 796, 9 to 13 min; [^13^C_6_]-UQ_9_ (M^+^ H^+^), *m/z* 801 to 802, 9 to 13 min. ECD and MS peak areas were corrected for sample loss during extraction using the recovery percentage of the internal standard UQ_10_. Quantification of UQ_9_, DQ_9_ and IDQ_9_ was performed using calibration curves generated from purified UQ_10_ standards, as previously described (32).

### Three-dimensional protein structure modeling and docking experiments

The 3D structure of UbiG from *P. aeruginosa* (protein ID, PA3171) complexed to SAM was predicted using AlphaFold 3 (33). The amino acid sequence of the target protein was submitted to the AlphaFold prediction pipeline, which uses multiple sequence alignments and structural templates, when available, to generate a 3D protein structure. The resulting model was evaluated using the predicted local distance difference test (pLDDT) score, which provides an estimate of the confidence of the prediction at the residue level. The predicted structure was subsequently visualized and analyzed using PyMOL. Regions displaying low prediction confidence were treated with caution and were not considered for detailed structural interpretation. Docking experiments were performed using Chai-1 prediction tool (34).

### Checkerboard synergy assay

Checkerboard assays were performed in triplicate, as previously described (35). Anaerobic cultures of the reference strain *P. aeruginosa* ATCC 27853 were grown in 96-well microplates containing the minimal medium described above, supplemented with two-fold serial dilutions of pABA and the antibiotic under investigation. The two compounds were arranged in a checkerboard format, with each well containing a unique combination of pABA and antibiotic concentrations. Following inoculation to a final density of approximately 10^5^ CFU/mL, plates were incubated for 20 h at 37 °C under anaerobic conditions. Bacterial growth was then assessed visually. The minimum inhibitory concentration (MIC) of each compound alone and in combination was determined as the lowest concentration preventing visible bacterial growth. The fractional inhibitory concentration index (FICI) of each combination was calculated as follows: FICI = MIC of pABA in combination / MIC of pABA alone (75 mg.L^-1^) + MIC of antibiotic in combination / MIC of antibiotic. The FICI values of the first well without growth along the turbid/non-turbid interface were averaged to determine interactions between compounds. Interactions were classified as synergistic when mean FICI ≤ 0.5, indifferent when 0.5 < mean FICI ≤ 4, and antagonistic when mean FICI > 4, as previously described (36).

## Data availability

The data underlying this article are provided as part of the main text and the supplementary data.

## Author contributions

L.P., J.M., M.L. and Y.C. designed the research and L.P. and M.L. obtained the fundings. L.P. performed all the bioinformatic analyses. J.M., O.L., T.A.D., and P.S. performed the experimental analyses and designed the corresponding figures. L.P. and J.M. wrote the original version of this manuscript with contributions from all co-authors. All authors agree with this version of the manuscript.

## Acknowledgments

This work was supported by the French National Research Agency and by the Grenoble Alpes University through the grants from the Agence Nationale de la Recherche, ANR-23-CE44-0015 (M.L.) and ANR-25-CE11-3606 (L.P.). The PhD of T.A.D was supported by a grant of the Graduate Schools/EUR from Grenoble Alpes University (ANR-17-EURE-0003).

## Abbreviations

CF: cystic fibrosis
UQ: ubiquinone
4-HB: 4-hydroxybenzoic acid
pABA: para-aminobenzoic acid
IDQ_9_: 4-imino-6-demethyl-UQ_9_
DQ_9_: 6-demethyl-UQ_9_
NPC: nonaprenylcatechol
NNPP: 2-amino-3-nonaprenylphenol.

## Conflict of interest

The authors declare no conflict of interest.

